# Genomic evidence that blind cavefishes are not wrecks of ancient life

**DOI:** 10.1101/2021.06.02.446701

**Authors:** Maxime Policarpo, Patrick Laurenti, Erik García-Machado, Cushla Metcalfe, Sylvie Rétaux, Didier Casane

## Abstract

Cavefishes often have modified eyes, from small but otherwise functional, to highly degenerate structures embedded in a connective tissue and covered by skin. Darwin assumed that these animals are ‘wrecks of ancient life’, but several genomic studies suggests they are not ‘ancient’. The most radical dating shift is for populations of a Mexican cavefish, *Astyanax mexicanus*, that have been recently estimated to be at the most a few tens of thousands years old. Despite having highly degenerate eyes, the eye-specific genes of *A. mexicanus* have low levels of decay. Other blind cavefishes we have examined so far are even older, but also can be dated to the Pleistocene. Here, we estimated the age of blindness of two additional fish species by the level of decay of eye-specific genes. Many pseudogenes were identified in the amblyopsid *Typhlichthys subterraneus*, suggesting that blindness evolved a few million years ago. In contrast, the blind cichlid *Lamprologus lethops* appears to be a new case of very recent and rapid eye regression, which occurred in deep river water, an environment similar to caves. Genome-wide analyses support the hypothesis that blindness in cavefishes is never very ancient, and ranges from the Early Pliocene to Late Pleistocene. Together with the description of hundreds of cavefish species, our results suggest that surface fishes were able to recurrently and rapidly adapt to caves and similar small dark ecosystems but the resulting highly specialized blind species with a limited distribution may be evolutionary dead-ends in a relatively short time.

## Introduction

In *On the origin of species*, Darwin wrote: “Far from feeling any surprise that some of the cave-animals should be very anomalous, as Agassiz has remarked in regard to the blind fish, the Amblyopsis, and as is the case with the blind Proteus with reference to the reptiles of Europe, I am only surprised that more wrecks of ancient life have not been preserved, owing to the less severe competition to which the inhabitants of these dark abodes will probably have been exposed” (Darwin 1859). Since the middle of the 19th century, thousands of obligate subterranean dwelling animals (troglobionts) have been described, most of them small arthropods (Culver and Pipan 2009; Romero 2009). Among the vertebrates, in addition to the cavefish, *Amblyopsis spelaea*, and the cave salamander, *Proteus anguinus*, mentioned by Darwin, many other troglobionts, primarily approximately two hundred and fifty named species of cavefishes (https://cavefishes.org.uk/), and at least 11 salamander species (Romero 2009) have been reported.

In the text quoted above, anomalous animals refers to troglomorphic (cave-related) characters, often present in cave animals, the two most conspicuous being highly degenerate eyes and depigmentation. Caves are very challenging ecosystems and surface animals have to adapt after settling in order to thrive in the long term. Interspecific competition is probably lower because there are fewer predators and other species that are competitors for food resources, but intraspecific competition is high in such a nutrient-limited environment (Poulson and White 1969). The depiction of cavefishes as wrecks of ancient life suggests that regressed traits are the result of relaxed selection, but it does not acknowledge that direct and indirect selection could also be involved in the evolutionary process. For instance, eye degeneration depends on mutations disrupting their development, but both selection and genetic drift could have been implicated in their fixation. In *Astyanax mexicanus* cavefish, a QTL analysis (Protas, et al. 2007) and indirect arguments based on the energetic cost of eyes (Moran, et al. 2015) suggest that selection for smaller eyes may have been involved in eye degeneration. Other analyses suggest indirect selection on eye size as the consequence of direct selection on other traits genetically correlated (Yamamoto, et al. 2009; Yoshizawa, et al. 2012; Hinaux, et al. 2016). In addition, if we assume that the first mutations reducing eye size were advantageous in the darkness of caves, other mutations may have accumulated later through genetic drift in small-eyed or blind fish (Retaux and Casane 2013). It has proven very difficult to disentangle these processes because they are not mutually exclusive (Retaux and Casane 2013). In addition, the term “wreck” does not acknowledge that cave animals in general, and cavefishes in particular, may evolve constructive traits involved in their adaptation to cave ecosystems (Hinaux, et al. 2016; Soares and Niemiller 2020).

Previous studies have used the level of eye regression to infer the relative age of cavefishes. However, these studies were biased and unreliable because they assumed that eye degeneration is a slow progressive process, lacked an explicit phylogenetic framework, or used phylogenies based on troglomorphic characters (Eigenmann 1909; Poulson and White 1969; Poulson 2001). Our current knowledge of eye development suggests that the idea of a slow, steady and progressive process is misleading because the impact of mutations on eye development can vary enormously, from a minor effect on eye structure and function to the most extreme case in which a single mutation can produce an eyeless phenotype (Loosli, et al. 2003). Today, phylogenetic inference and dating based on gene sequences have superseded those relying on morphological characters. The main advantage of a gene-based phylogenetic approach is that relationship and dating inferences are independent of the analysis of the evolution of morphological traits which are mapped on *a priori* molecular phylogenies. Assuming that genes used for phylogenetic analyses and troglomorphic traits evolved independently, molecular phylogenies allow us to identify convergent evolution. Moreover, the analysis of genes considered to be under relaxed purifying selection after cave settlement allows us to date the environmental shift from surface to cave and to test hypotheses on the mode and tempo of eye degeneration.

A recent study of the genomes of several cavefish species available at the time allowed us to examine the decay of many eye-specific genes and non-visual opsins, and estimate the age of the shift from surface to cave environment (Policarpo, et al. 2021). This genome-wide analysis demonstrated a low level of gene decay suggesting that these cavefishes originated recently, that is during the Pleistocene (from 2,580,000 to 11,700 years ago). Moreover, the level of eye regression was not correlated with the proportion of eye-specific pseudogenes. However, previous studies based on analyses of one or two genes suggest that other cavefishes could be much more ancient, in particular some among the amblyopsid cavefishes (Dillman, et al. 2011; Niemiller, et al. 2013). Here we sought to further investigate the time of origin of a blind amblyopsid cavefish, *Typhlichthys subterraneus*, using our genome-wide approach. We also examined the same set of eye-specific genes and non-visual opsins of a blind cichlid, *Lamprologus lethops*, which inhabits the deep-water of the Congo river. The level of gene decay in *Typhlichthys subterraneus* supports that blindness may be a few million years old in this lineage, indicating that this species may had settled in caves during the Pliocene (from 5.333 million to 2.58 million years ago). In contrast, the very low level of gene decay in *Lamprologus lethops* indicates that eye regression is very recent, that is it evolved less than 100,000 years ago. These analyses strengthen the hypothesis that once fishes have settled into a dark environment, eyes can degenerate rapidly, reducing in size, sinking into their orbit and becoming covered by skin. In parallel, genes with eye-restricted expression accumulate LoF mutations, but this process is comparatively much slower.

## Results

### Identification of LoF mutations and pseudogenes in vision, circadian clock and pigmentation genes

In a previous study (Policarpo, et al. 2021), we defined three gene sets based on the zebrafish genome: 95 vision genes, 42 circadian clock genes and 257 pigmentation genes (**fig. 1**). In the present analysis, we looked for likely LoF mutations (i.e. premature STOP codons, losses of START and STOP codons, losses of intron splice sites and small indels leading to frameshifts) in homologs from the genomes of a blind cavefish from North America, the Southern Cavefish *Typhlichthys subterraneus* (Amblyopsidae) and a blind fish living in deep water of the Congo River in Central Africa, *Lamprologus lethops* (Cichlidae). For each species, *T. subterraneus* and *L. lethops*, we also examined the genes in the closest related surface fish for which a genome was available, *Percopsis transmontana* and *Lamprologus tigripictilis*, respectively, and a more distantly related surface species, *Gadus morhua* and *Neolamprologus pulcher*, respectively (**fig. 2**).

**Fig 1:**
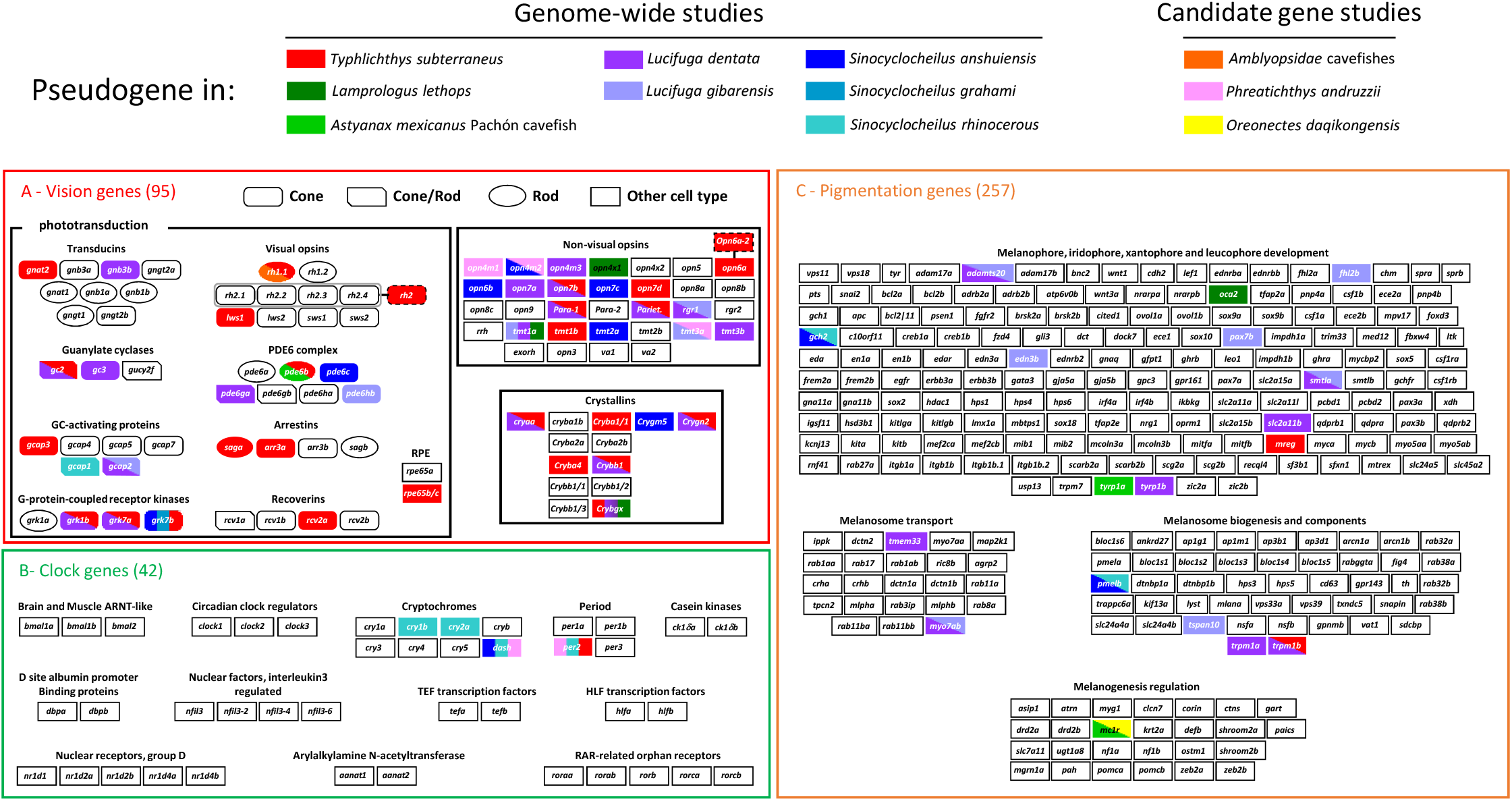
Gene sets and pseudogenes. (A) Vision genes. (B) Circadian clock genes. (C) Pigmentation genes. Pseudogenes are coloured according the species in which they were found. In candidate gene studies, only few genes were examined, whereas most genes were inspected in our genome-wide analyses. Gene copy number variations: there are four copies of *Rh2* in zebrafish whereas there is only one *Rh2* gene in *T. subterraneus*; there is one *opn6a* gene in zebrafish whereas there are two *opn6a* genes in *T. subterraneus*. Data for previously studied cavefishes (*A. mexicanus, Lucifuga* spp. and *Sinocyclocheilus* spp.) have been redrawn from Policarpo et al. (2021), for comparison.

**Fig 2:**
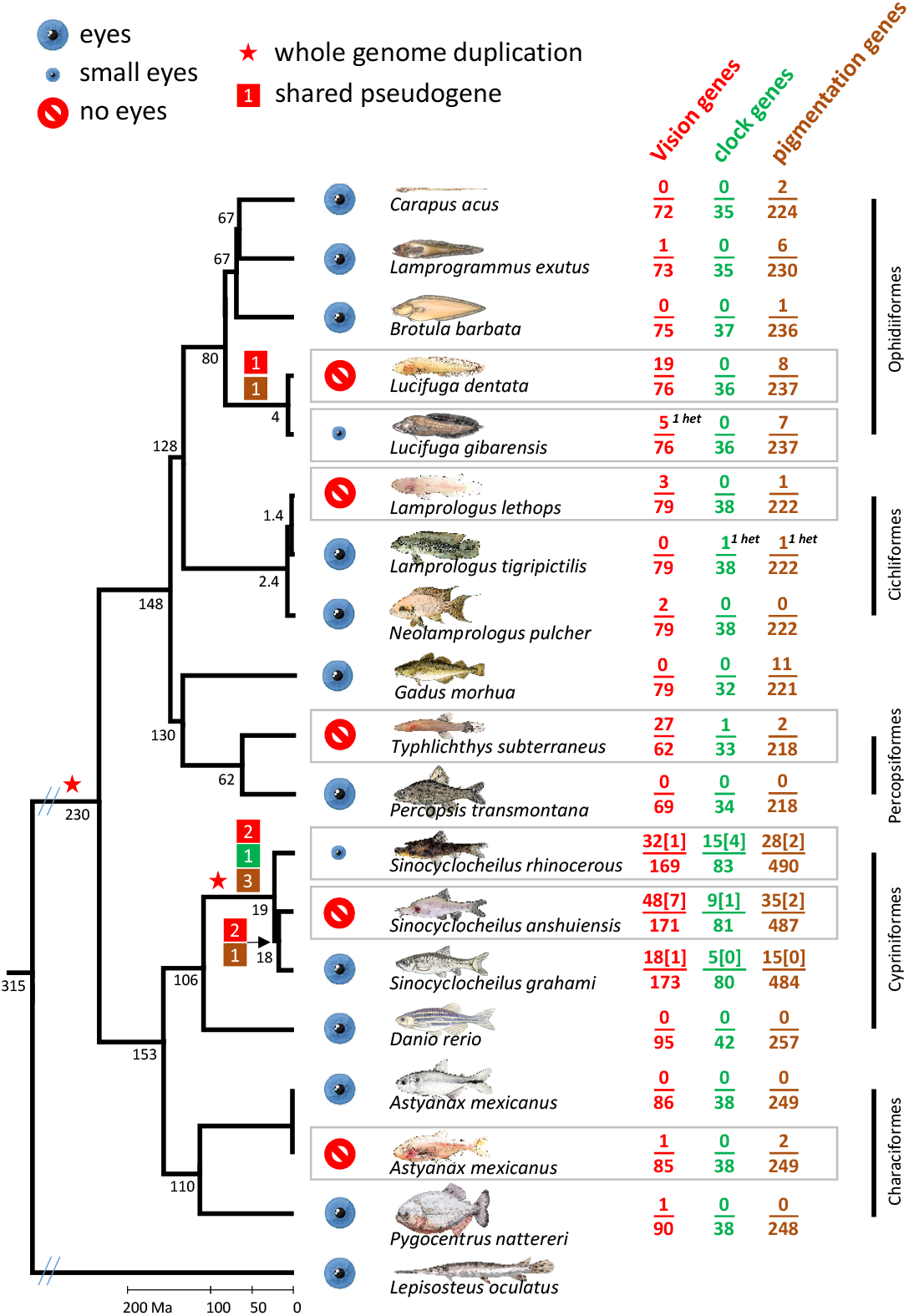
Phylogeny and pseudogene mapping. For each gene set, the number of pseudogenes found and the number of gene examined in a species are given to the right of the species name. het: heterozygous. Data for previously studied cavefishes (*A. mexicanus*, *Lucifuga* spp. and *Sinocyclocheilus* spp.) have been redrawn from Policarpo et al. (2021), for comparison.

A total of 55 LoF mutations were found in *T. subterraneus* and *L. lethops*. Most LoF mutations (47) were found in vision genes, while only 2 LoF were found in circadian clock genes and 6 in pigmentation genes. Most LoF mutations in vision genes (43/47) were found in *T. subterraneus* (**supplementary fig. S1 and fig. S2**, **Supplementary Material** online). In order to test if these LoF mutations were distributed randomly within the genes, that is, they were not clustered at the 3′ end of the genes where their deleterious effect would be smaller, we computed the effective number of gene segments generated by LoF mutations and compared this value with simulations of random distributions of mutations within genes (Policarpo, et al. 2021). We found that premature STOP codons and frameshifts are distributed randomly throughout the coding sequences (**supplementary fig. S3**, **Supplementary Material** online).

We counted the number of pseudogenes in all six species, that is, the number of genes carrying at least one LoF mutation (**fig. 1** and **fig. 2**). Among the vision genes, we identified 27 pseudogenes in *T. subterraneus*, but only 3 pseudogenes in *L. lethops*. The LoF mutations found in *L. lethops* (**fig. 1**, **supplementary fig. S2**, **Supplementary Material** online) have been reported in a previous study (Aardema, et al. 2020). A gene, *opn3*, was also reported as pseudogenized in this publication, but it was considered a functional gene in our study. Indeed, the frameshift reported in *opn3* is probably an artefact as this gene is not where the deletion, at the origin of the frameshift, was found. We did not find any LoF mutations in this gene (for more details, see **supplementary fig. S4**, **Supplementary Material** online).

The percentage of vision pseudogenes in *T. subterraneus* (44%) is the highest ever reported in cavefishes. In contrast, the percentage of vision pseudogenes is very low in *L. lethops* (4%). No circadian clock pseudogene was found in *L. lethops* and only one pseudogene (*per2*) in *T. subterraneus*. The loss of *per2* is recurrent in cavefishes, as pseudogenes of *per2* have been identified in *S. rhinocerous* and *P. andruzzii* (Yang, et al. 2016; Ceinos, et al. 2018; Policarpo, et al. 2021). A circadian clock pseudogene (*cry-dash*) was found in the surface fish *L. tigripictilis*, however, the specimen was heterozygous for this gene. Two pigmentation pseudogenes (*trpm1b* and *mreg*) were identified in *T. subterraneus* and one pseudogene (*oca2*) in *L. lethops*. This mutation has already been reported in a previous study (Aardema, et al. 2020). In the surface fish *L. tigripictilis*, a pseudogene (*pts*) was found, however, the specimen was also heterozygous for the gene (**fig. 2**, **Table S1**, **Supplementary Material** online).

Together, findings from the present study and a previous study (Policarpo, et al. 2021) has allowed us to estimate that among the set of 95 vision genes, 46 have been found pseudogenized in at least one cave species, that is, 48% of the gene set. In sharp contrast, only 4 pseudogenes were identified among 42 circadian clock genes, and 18 pseudogenes among the 257 pigmentation genes, that is, 10% and 7% respectively.

### Estimation of the number of neutral vision genes based on the distribution of LoF mutations per pseudogene

In *T. subterraneus*, among 62 vision genes, 43 LoF mutations were found in 27 pseudogenes. More precisely, there were 17 pseudogenes with 1 mutation, 7 with 2 mutations, 1 with 3 mutations, 1 with 4 mutations and 1 with 5 mutations. This distribution was compared with expected distributions obtained for different numbers of neutral genes ranging from 27 (44%) to 62 (100%). A better fit between the observed and expected distribution was found when at least 80% of genes were assumed to be neutral sequences (**fig. 3**). This result is congruent with the estimation based on the distribution of LoF mutations per pseudogene in *Lucifuga dentata* (Policarpo, et al. 2021) (**fig. 3**). Moreover, the higher percentage of pseudogenes with more than one LoF mutation in *T subterraneus* than in *L. dentata* is congruent with the higher percentage of pseudogenes (44% *vs* 25%) indicating that *T. subterraneus* is the oldest cavefish species.

**Fig 3:**
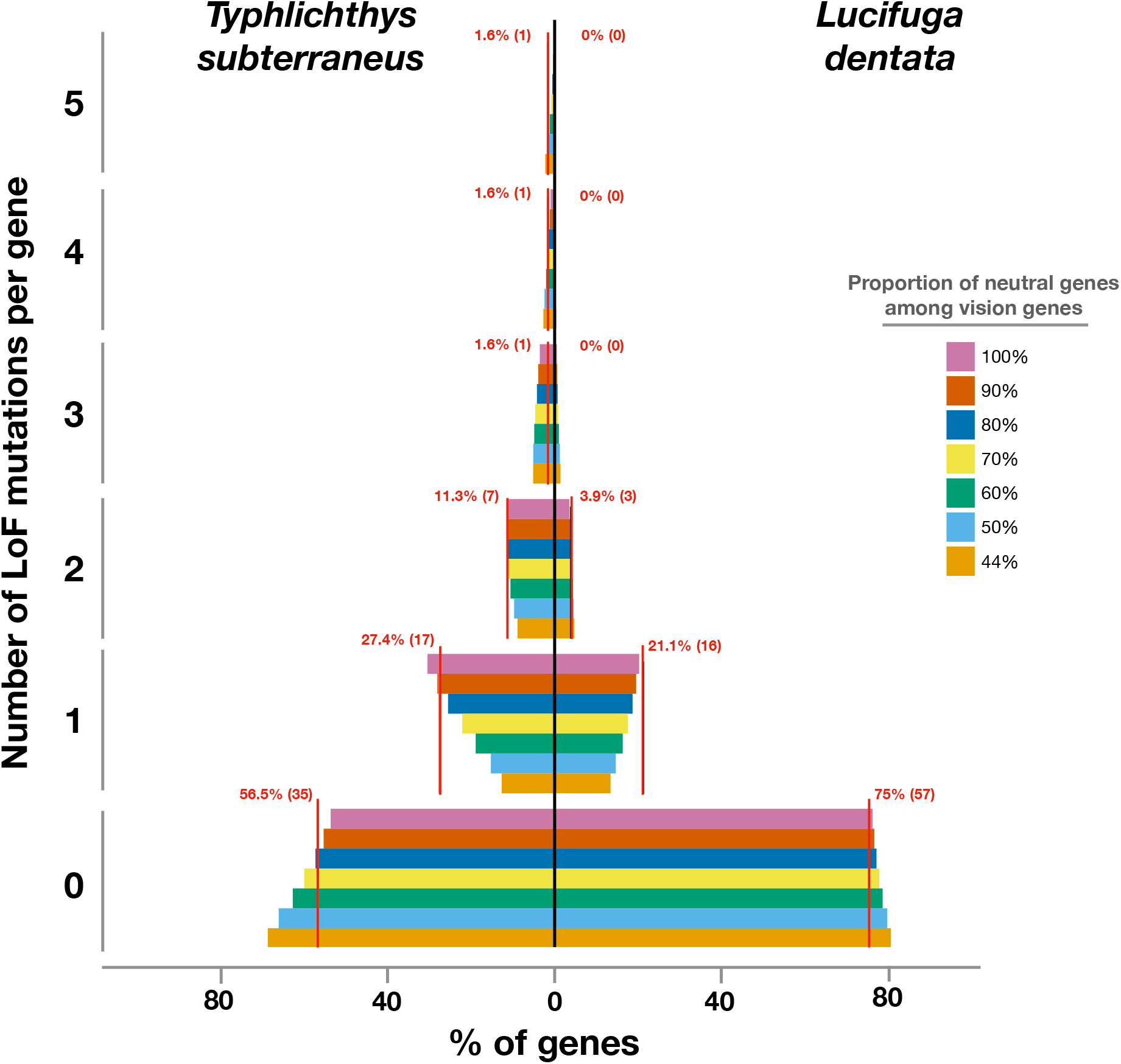
Estimation of the number of neutral genes in *T. subterraneus*. Red lines: observed distribution. Expected distributions were obtained for different numbers of neutral genes among 62 genes: between 27 and 62 (each expected distribution was obtained with 10,000 simulations). The distributions for *L. dentata* have been redrawn from Policarpo et al. (2021), for comparison.

Another approach to tackle this issue involved the analysis of the number of vision genes pseudogenized in both species. There are 27 pseudogenes among 62 vision genes in *T. subterraneus* (present study) and 19 pseudogenes among 76 vision genes in *L. dentata* (Policarpo, et al. 2021). Moreover, among 58 vision genes identified in both species, there are 10 genes - *opn7b, parapinopsin-1, parietopsin, cryaa, crybb1, crybgx, crygn2, grk1b, grk7a* and *gc2* - pseudogenized in both species. As expected for two distantly related blind species, no shared LoF mutation was found in orthologous pseudogenes (**supplementary Table S2**, **Supplementary Material** online). Assuming that all vision genes are dispensable and have the same LoF mutation rate, we can compute the expected number of genes pseudogenized in parallel in two species among a set of vision genes studied in both species (probability of pseudogenization in one species x probability of pseudogenization in the other species x number of genes), that is 27/62 × 19/76 × 58 = 6.3. A script was written to compute the distribution of the probabilities of the number of shared pseudogenes, in a range between 0 and 14, taking into account that 14 pseudogenes were found in *L. dentata* and 25 in *T. subterraneus* among the 58 vision genes found in both species (**supplementary fig. S5**, **Supplementary Material** online). The expected value (6.3) and the low probability (0.01) of finding 10 shared pseudogenes suggest a significant excess of shared pseudogenes that could be evidence that some vision genes cannot carry LoF mutations. However, with a more realistic model taking into account the length and the number of introns of each gene to estimate the relative LoF mutation rate of each gene, a probability higher than 5% was found for a number of shared pseudogenes in a range between 5 and 10 (**supplementary fig. S5**, **Supplementary Material** online). This result is another evidence that most, if not all, vision genes are dispensable in these blind species.

### Dating the relaxation of purifying selection on vision genes in four blind fishes

To date the relaxation of purifying selection on vision genes in *T. subterranues, L. lethops* and *A. mexicanus*, for each species, we used the number of pseudogenes among vision genes and the LoF mutation rate per generation estimated as described in a previous publication (Policarpo, et al. 2021). The highest probability of finding 27 pseudogenes among 62 genes in *T. subterraneus* was obtained for relaxed selection starting 569,570 generations ago (probability > 5% between 447,390 and 712,830 generations). For *L. lethops* (3 pseudogenes among 79 genes), the time since relaxed selection was estimated to 28,970 generations ago (probability > 5% between 8,670 and 68,620 generations). For *A. mexicanus* (1 pseudogene among 85 genes), the time since relaxed selection was estimated to 8,560 generations ago (probability > 5% between 450 and 38,580 generations). For *L. dentata* (19 pseudogenes among 76 genes), the estimate was obtained in a previous study (Policarpo, et al. 2021), 367,780 generations ago (probability > 5% in a range between 273,990 and 480,980 generations)(**fig. 4**).

**Fig 4:**
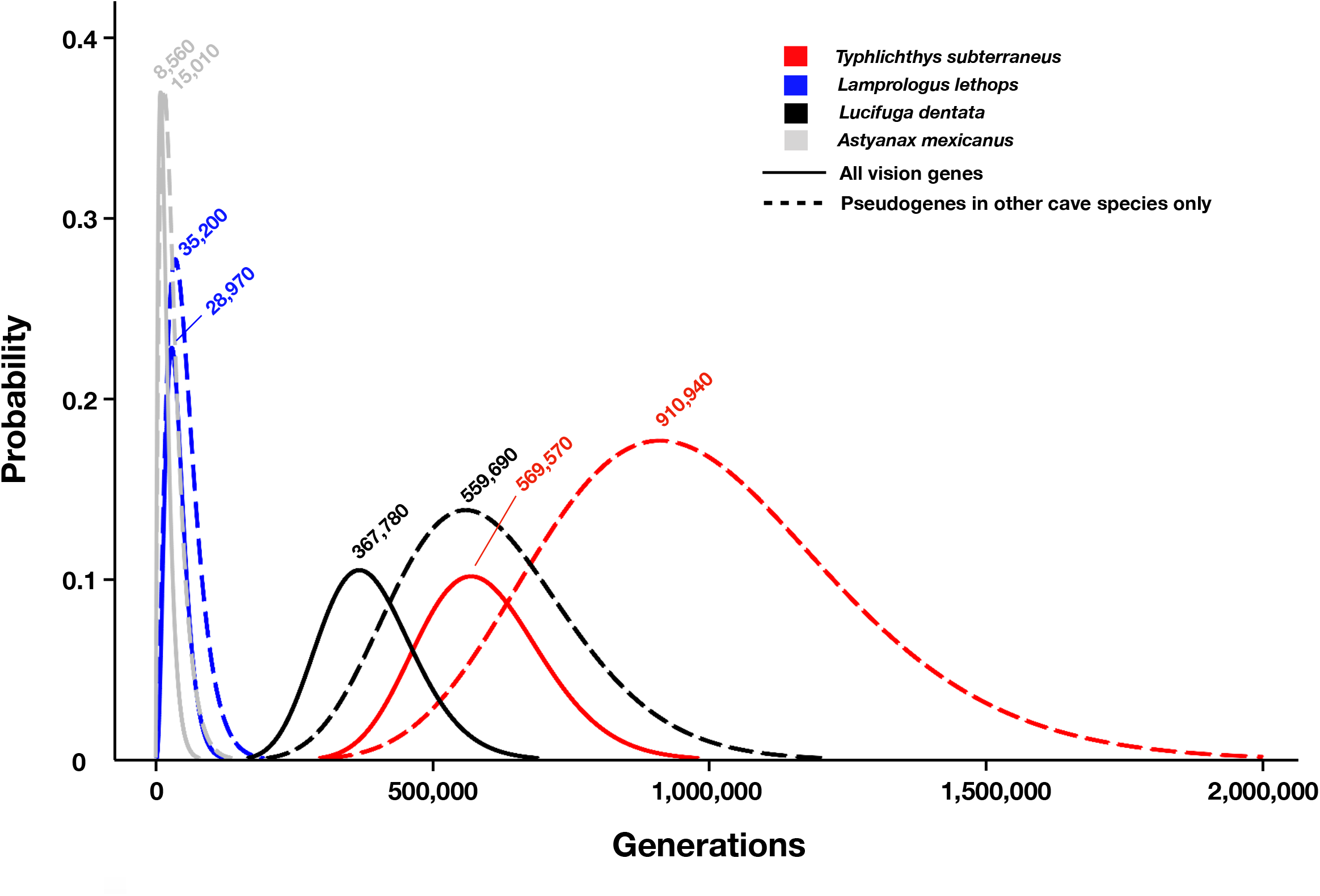
Dating of the relaxation of purifying selection on vision genes. Solid lines: probabilities of finding 27 pseudogenes among 62 vision genes in *T. subterraneus*, 19 pseudogenes among 76 genes in *L. dentata*, three pseudogenes among 79 vision genes in *L. lethops* and one pseudogene among 85 vision genes in *A. mexicanus*, according to the time in caves (generations) and assuming relaxed selection on the whole vision gene set in this environment. Dashed lines: probabilities of finding pseudogenes using only genes found pseudogenized in other cave species, increasing the likelihood that the genes examined are dispensable in caves. The number of generations for which the highest probability was found is reported.

For each species, we also computed this probability using only a subset of vision genes found pseudogenized in at least one among the other cavefishes, increasing the likelihood that the genes examined are dispensable in caves. Excluding *T. subterraneus*, 33 genes were found with at least one LoF mutation in the other cave species. Twenty-one of these genes were identified in *T. subterraneus* of which 13 were pseudogenes. The highest probability of finding 13 pseudogenes among 21 genes was obtained for relaxed selection starting 910,940 generations ago (probability > 5% between 557,730 and 1,401,750 generations). Excluding *L. lethops*, 46 genes with a LoF were found in the other cave species. Forty-one of these were identified in *L. lethops*, of which 2 were pseudogenes. The time since relaxed selection was estimated to be 35,200 generations ago (probability > 5% between 6,640 and 103,250 generations). Excluding *A. mexicanus*, 47 genes with a LoF were found in the other cave species. Forty-four of these were identified in *A. mexicanus*, of which one was a pseudogene. The time since relaxed selection was estimated to be 15,010 generations ago (probability > 5% between 790 and 67,740 generations). Excluding *L. dentata*, 41 genes with a LoF were found in the other cave species. Thirty-five of these were identified in *L. dentata*, of which 13 were pseudogenes. The time since relaxed selection was estimated to be 559,690 generations ago (probability > 5% between 365,220 and 814,710 generations).

## Discussion

We discuss the patterns of gene decay among different gene sets in various cavefishes. Then, we show how our analyses of the genomes of a blind cavefish and a deep-water blind fish strengthen our hypotheses on the pace of eye loss and the evolutionary fate of cave-adapted fishes.

### Patterns of gene decay

In a previous study, we defined a set of genes we called vision genes that are most likely to be dispensable in a dark environment and can carry LoF mutations without a large impact on overall fitness. They are primarily eye-specific genes involved in visual phototransduction, but we also included a set of non-visual opsin that showed similar gene decay (Policarpo, et al. 2021). In this publication, our analysis of gene decay in *L. dentata* suggested that most, if not all, vision genes are dispensable in blind cave fishes, however only 25% of *L. dentata* vision genes have LoF mutations. In the present study, we found that 44% of the vision genes of *T. subterraneus* have LoF mutations, an observation strongly strengthening our hypothesis. In *T. subterraneus*, the distribution of LoF mutations per genes is similar to the one in *L. dentata* and in accordance with the assumption that most vision genes are under relaxed purifying selection in the darkness of caves.

The analysis of seven cavefishes identified pseudogenes in 48% of the 95 vision genes. We have shown that this set of genes is a useful tool for estimating the time of origin of blindness and the extent of common origin and parallel evolution among closely related cavefishes. However, only the analysis of other cavefish genomes may allow us to unambiguously identify the subset of vision genes that are dispensable in the dark.

### Decoupling of eye regression and vision gene decay

Assuming that eye regression is a progressive process and that cavefishes can be ordered on a ladder of degeneration, eye regression has been used to estimate the age of different cavefish lineages (Wilkens, et al. 1989). However, a well-established and independent phylogenetic framework and independent evidence of the age of eye regression are necessary to understand the dynamic of eye regression and the diversity of its modality in different cavefish lineages. The very low decay in vision genes in *Astyanax mexicanus* cavefish, despite their highly degenerate eyes, indicates that *A. mexicanus* has lost eyesight very recently (Fumey, et al. 2018; Policarpo, et al. 2021). The very similar pattern of gene decay observed in *Lamprologus lethops* suggests that eyesight was also lost very recently in this species, however, while the eyes are very small and sunk under the skin, their general organization is much better preserved. On the other hand, both cavefishes that have accumulated many vision pseudogenes, that is *Lucifuga dentata* and *Typhlichthys subterraneus*, have highly degenerate eyes. There are probably many evolutionary pathways leading to eye regression in cavefishes, some through intermediary stages involving first eye size reduction but normal development, while in other cases through changes in early development leading to highly degenerate eyes. Both chance and environmental constraints may be involved in this process. In particular, it is possible that some blind fishes need to retain the capacity to detect sources of light.

### Conclusion

The analysis of two recently published genomes of blind fishes, one living in caves and the other in the deep-water of a river, strengthens our hypothesis that blind cavefishes are not very ancient and most of them lost eye function during the Pliocene and Pleistocene. The recent eye regression in *L. lethops* indicates that eyes can sink rapidly without other important morphological changes, or as in *A. mexicanus*, with extensive eye degeneration. In both cases, the number of eye-specific pseudogenes is very small, two and one respectively. In other words, eyes can sink rapidly but eye-specific genes rust slowly in cavefishes.

Among blind cavefishes for which the genome is available, several lines of evidence suggest that the oldest may be the amblyopsid *T. subterraneus*, in which 44% of vision genes are pseudogenized. Nevertheless, other North American amblyopsids may be older (Niemiller, et al. 2013). When the genomes of these species will be available, the analysis of the vision genes should clarify the age of these cavefishes and also to which extent cave amblyopsids evolved independently.

The analysis of the genome of *T. subterraneus* allowed us to identify a blind cavefish that is probably older than *L. dentata*, which was previously considered the oldest cavefish among those with sequenced genomes, but this does not refute our current working hypothesis that blind cavefishes cannot thrive more than a few millions years in cave ecosystems.

## Materials and Methods

### Vision, circadian clock and pigmentation gene sets

The three sets of genes, vision, circadian clock and pigmentation, have been previously described (Policarpo, et al. 2021). Briefly, the set of vision genes was based on the expression patterns of 95 zebrafish genes coding for visual and non-visual opsins, crystallins and proteins involved in the phototransduction cascade. The set of circadian clock genes (42 genes in zebrafish) and the set of pigmentation genes (257 genes in zebrafish) were established in other studies (Li, et al. 2013; Lorin, et al. 2018). The complete list of genes with standard identifiers (https://zfin.org/) is given in **fig. 1**.

### Construction of data sets

*Danio rerio* sequences were used as queries for Blast and tBlastn 2.6.0+ (Altschul, et al. 1990) to the genome of *Percopsis transmontana* (GCA_900302285.1), *Gadus morhua* (gadMor1), *Neolamprologus pulcher* (GCF_000239395.1) and *Typhlichthys subterraneus* (GCA_900302405.1). Regions corresponding to the best matches were extracted with SAMtools v1.9 faidx (Li 2011). Coding sequences and introns were predicted using EXONERATE v2.4.0 (Slater and Birney 2005) (model protein2genome, using *Danio rerio* protein sequences as query).

BAM files of *Lamprologus lethops* and *Lamprologus tigripictilis* reads aligned to the genome of *Neolamprologus pulcher* were downloaded from NCBI (SRR10811633, SRR10811634). SAMtools BCFtools v1.9 (Li 2011) was used to generate a consensus genome (BCFtools consensus) from which genes were predicted following the method described above. Coding sequences were classified as “complete” if the complete CDS was retrieved or “incomplete” otherwise. If less than 10% of a coding sequence was found, the gene was classified as “Not Found” and it was not further analyzed. A CDS was classified as a “pseudogene” if at least one LoF mutation was found (premature STOP codon, loss of the initiation codon, loss of the STOP codon, indel leading to a frameshift, mutations at intron splice sites).

### Phylogenetic analyses

Orthologous and paralogous relationships between genes were inferred through phylogenetic analyses. Coding sequences were trimmed and aligned with MACSE v2.03 program trimNonHomologousFragments and alignSequences (Ranwez, et al. 2018). Frameshifts and internal stop codons were then removed with the MACSE v2.03 progam exportAlignment. For each alignment, DNA sequences were translated into protein sequences and a maximum likelihood phylogenetic tree was inferred using IQ-TREE v1.6.12 (Nguyen, et al. 2015) with the optimal model found by ModelFinder (Kalyaanamoorthy, et al. 2017). Robustness of the nodes was evaluated with 1,000 ultrafast bootstraps (Hoang, et al. 2018). Trees were rooted and visualized using iTOL v6 (Letunic and Bork 2007).

### Estimation of the proportion of neutral vision genes in cavefishes

The distribution of LoF mutations per gene in *T. subterraneus* was used to estimate the proportion of neutral genes using the method described previously in (Policarpo, et al. 2021). In brief, 10,000 simulations of the distribution of 43 mutations in a random sample of V genes from the 62 *T. subterraneus* vision genes was generated. V ranged from the number of observed pseudogenes (27), which is the minimal number of neutral genes, to the total number of genes (62). The length of the coding sequence and the number of introns of each gene were taken into account to compute the relative mutation rate.

Then, we computed the expected number of vision genes pseudogenized in both *T. subterraneus* and *L. dentata*, assuming that all vision genes are dispensable and have the same LoF mutation rate. The following formula was used: E = P_Ts_ x P_Ld_ x N, where P_Ts_ is the probability that a vision gene is pseudogenized in *T. subterraneus*, P_Ld_ is the probability that a vision gene is pseudogenized in *L. dentata*, and N is the number of vision genes found in both species.

Given that of the 58 vision genes found in both species, 14 genes were pseudogenized in *L. dentata*, 25 in *T. subterraneus*, and 10 were shared, a custom R script was used to evaluate if this number of shared pseudogenes was likely under the hypothesis that all vision genes are dispensable. The script generated 10,000 simulations of random pseudogenization of the genes, taking into account the length and the number of introns of each gene to compute its relative mutation rate. For each simulation, the number of shared pseudogenes was recorded, allowing us to estimate the probability of obtaining a given number of shared pseudogenes in the range between 0 and 14 (**supplementary fig. S5**, **Supplementary Material** online).

### Dating relaxation of purifying selection on vision genes in *L. lethops* and *T. subterraneus* from pseudogene numbers

We used the method previously described (Policarpo, et al. 2021). Briefly, we computed the probability than D genes among T genes accumulated a LoF mutation taking into account the mean gene length. This method requires an estimate of the LoF mutation rate per generation. Assuming a transition/transversion ratio in vision genes of 1.68 computed with PAML v4.9h (Yang 2007) and taking into account the codon usage in vision genes, we found that the probability that a single mutation led to a premature stop codon was 0.031. In *T. subterraneus*, 12 premature stop codons and 27 frameshifts were found in vision genes. Thus, the probability that a mutation is a frameshift was (27/12) × 0.031 = 0.069.

The probability of splice site mutation was 4 x (total intron number / total cds length) = 4 x (255/48703) = 0.021. The probability of start loss was equal to 3 x (number of genes / total cds length) = 3 x (62/48703) = 0.0038 and the probability of stop loss was equal to 0.852*(Start loss probability) = 0.0033. Thus μ_LoF_ was equal to 0.128μ with μ set to 10^−8^. As a very low number of pseudogenes were found in *L. lethops*, the LoF mutation rate found in *T. subterraneus* was also used for dating relaxation of selection in *L. lethops and A. mexicanus*. Dating of *L. dentata* was estimated using the LoF mutation rate (0.0717μ) reported in a previous study (Policarpo, et al. 2021).

### R analyses

R script files for reproducing analyses and figures is available in supplementary files (see Data Availability) and on GitHub (https://github.com/MaximePolicarpo/Genomic-evidence-that-cavefishes-are-not-wrecks-of-ancient-life). A list of R packages used is given below, with a reference to a publication if available or an URL otherwise: plotrix v3.7-7 (Lemon 2006), data.table v1.12.8 (https://cran.r-project.org/web/packages/data.table/index.html), dplyr v0.8.5 (https://cran.r-project.org/web/packages/dplyr/index.html), tidyverse v1.3.0 (Wickham, et al. 2019), purr v0.3.3 https://cran.r-project.org/web/packages/purrr/index.html, ggplot2 v3.3.0 (Wickham 2016), ggpubr v0.2.5 (https://cran.r-project.org/web/packages/ggpubr/index.html), gtools v3.8.2 (https://cran.r-project.org/web/packages/gtools/index.html), forcats v0.5.0 (https://cran.r-project.org/web/packages/forcats/index.html), FSA v0.8.30 (Ogle, et al. 2020), patchwork v1.0.0 (https://cran.r-project.org/web/packages/patchwork/index.html), quantable 0.3.6 (https://cran.r-project.org/web/packages/quantable/index.html), reticulate v1.15 (https://cran.r-project.org/web/packages/reticulate/index.html), sjPlot v2.8.3 (Lüdecke 2020), gridExtra v2.3 (https://cran.r-project.org/web/packages/gridExtra/index.html), ape v5.3 (Paradis and Schliep 2019) and phytools v0.7-47 (Revell 2012).

## Data Availability

All data and results of the analyses performed in this study are available for download at figshare (https://figshare.com/articles/dataset/Supplementary_files/14237279): three tables with all gene sequences and their genomic positions (one table per gene data set); three tables with a description of LoF mutations (one table per gene data set); Exonerate raw results; Gene alignments, IQ-TREE results and phylogenetic trees in PDF; a R script to reproduce the analyses.

## Supplementary Material

Supplementary data are available at Genome Biology and Evolution.

## Acknowledgments

This work was supported by a collaborative grant from Agence Nationale de la Recherche (BLINDTEST to S.R. and D.C.) and from Institut Diversité Ecologie et Evolution du Vivant (to S.R. and D.C).

