## Supplementary Material - Tables and Figures for "Genomic evidence that blind cavefishes are not wrecks of ancient life"

**Table S1.** Pseudogene read counts in *Lamprologus* spp..

**Table S2.** Number of vision genes, pseudogenes and LoF mutations retrieved from *L. dentata* and *T. subterraneus*.

**Fig. S1.** A - Distribution of the different types of LoF mutations in *T. subterraneus* and *L. lethops*. B - Distribution of frameshift sizes.

**Fig. S2.** A - List of vision genes retrieved from cavefishes and related species. Colors represent the LoF mutation type. When higher than one, the number of LoF mutation is also reported. B - Circadian clock pseudogenes retrieved from cavefishes and related species. C - Pigmentation pseudogenes retrieved from cavefishes and related species.

**Fig. S3.** Distribution of the effective segment size generated by random insertion of STOP codons and frameshifts (100,000 simulations).

**Fig. S4.** Visualization of the artefactual loss-of-function mutation found by Aardema *et al.* (2020) in *opn3* of *L. lethops*. A – No gene found in the *N. brichardi* genome at the location given in Aardema *et al.* (2020) showing no gene annotated in this region. B - Exonerate result and proper location of *opn3* on the *N. brichardi* genome. C - Protein alignment of *opn3* of various teleost species showing that *Lamprologus lethops* sequence is complete and without LoF mutations.

**Fig. S5.** Probability of finding a number of shared pseudogenes among 58 genes found in both *L. dentata* and *T. subterraneus*.

Table. S1

| **Species** | **Gene** | **Scaffold** | **Position** | **DP4 field** | **Genotype** |
|---|---|---|---|---|---|
| *Lamprologus lethops* | crybgx | NW_006272218.1 | 642367 | 1,5,23,26 | 1/1 |
| *Lamprologus lethops* | oca2 | NW_006272018.1 | 5745343 | 0,0,25,26 | 1/1 |
| *Lamprologus lethops* | opn4x1 | NW_006272015.1 | 5259668 | 0,0,26,50 | 1/1 |
| *Lamprologus lethops* | tmt1a | NW_006272087.1 | 1747364 | 3,0,30,15 | 1/1 |
| *Lamprologus lethops* | tmt1a | NW_006272087.1 | 1747251 | 5,7,11,16 | 0/1 |
| *Lamprologus tigripictilis* | cry-dash | NW_006272036.1 | 5029708 | 7,9,32,10 | 0/1 |
| *Lamprologus tigripictilis* | pts | NW_006272048.1 | 1127316 | 18,13,14,16 | 0/1 |

Table. S2

| Species | Number of vision genes retrieved | Number of vision genes retrieved with at least 1 LoF mutation | Number of LoF mutations retrieved in vision genes |
|---|---|---|---|
| *L. dentata* | 76 | 19 | 22 |
| *T. subterraneus* | 62 | 27 | 43 |
| Common | 58 | 10 | 0 |

Fig. S1

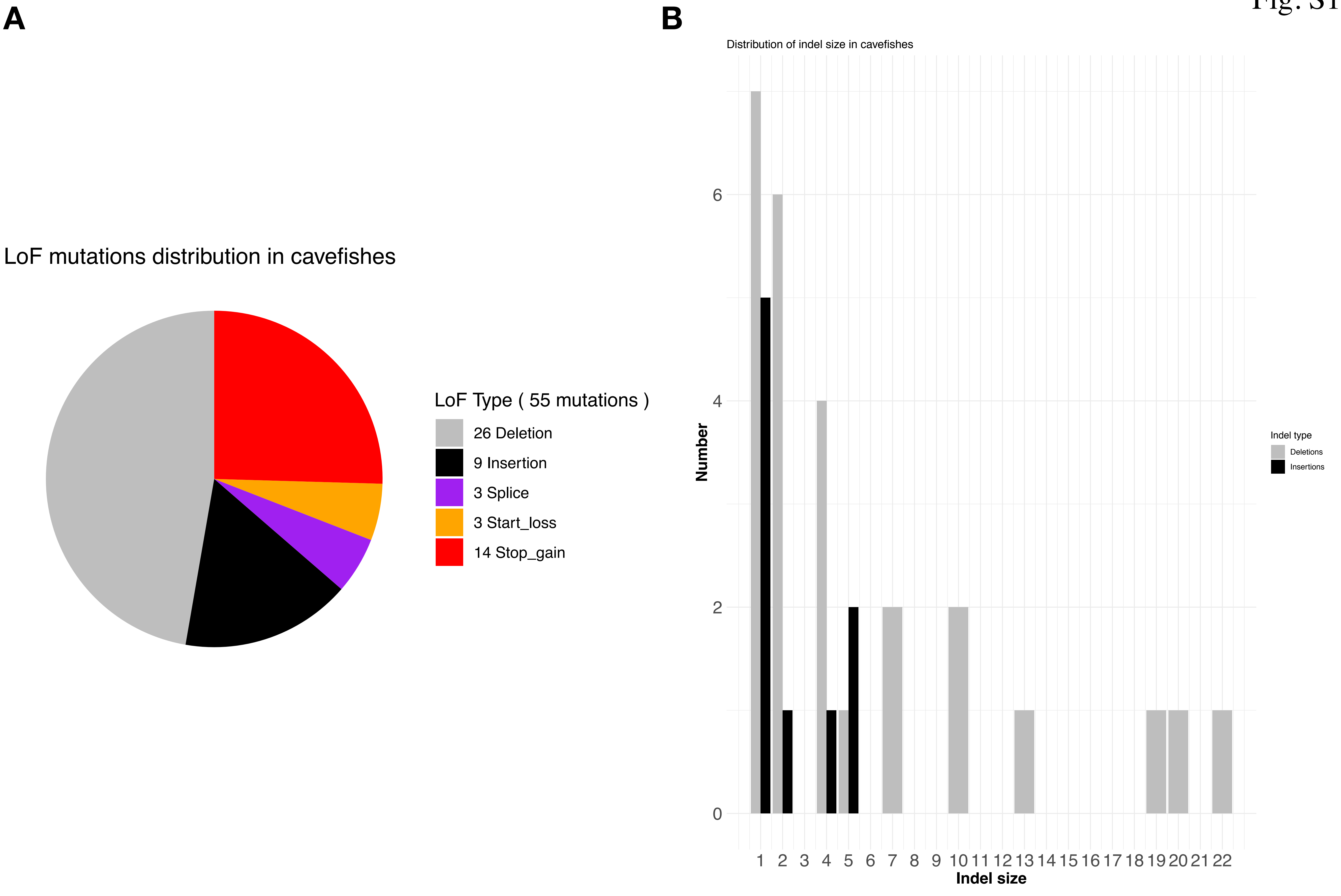

A
LoF mutations distribution in cavefishes
LoF Type ( 55 mutations )
26 Deletion
9 Insertion
3 Splice
3 Start_loss
14 Stop_gain
B
Distribution of indel size in cavefishes
Number
0
2
4
6
1 2 3 4 5 6 7 8 9 10 11 12 13 14 15 16 17 18 19 20 21 22
Indel size
Indel type
Deletions
Insertions

Fig. S2

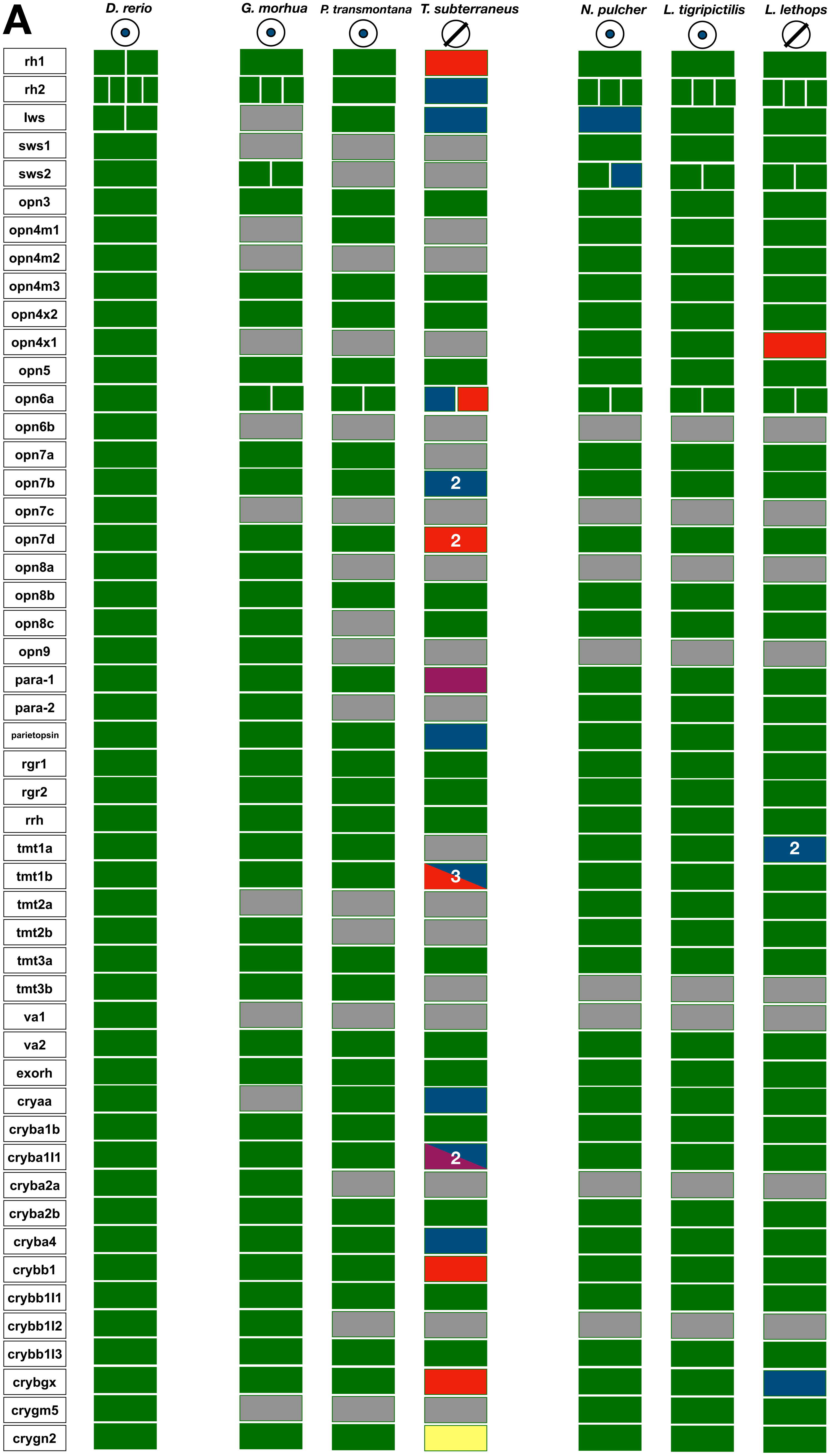
A
D. rerio
G. morhua
P. transmontana
T. subterraneus
N. pulcher
L. tigripictilis
L. lethops
rh1
rh2
lws
sws1
sws2
opn3
opn4m1
opn4m2
opn4m3
opn4x2
opn4x1
opn5
opn6a
opn6b
opn7a
opn7b
opn7c
opn7d
opn8a
opn8b
opn8c
opn9
para-1
para-2
parietopsin
rgr1
rgr2
rrh
tmt1a
tmt1b
tmt2a
tmt2b
tmt3a
tmt3b
va1
va2
exorh
cryaa
cryba1b
cryba1l1
cryba2a
cryba2b
cryba4
crybb1
crybb1l1
crybb1l2
crybb1l3
crybgx
crygm5
crygn2
2
2
2
3
2

Fig. S2

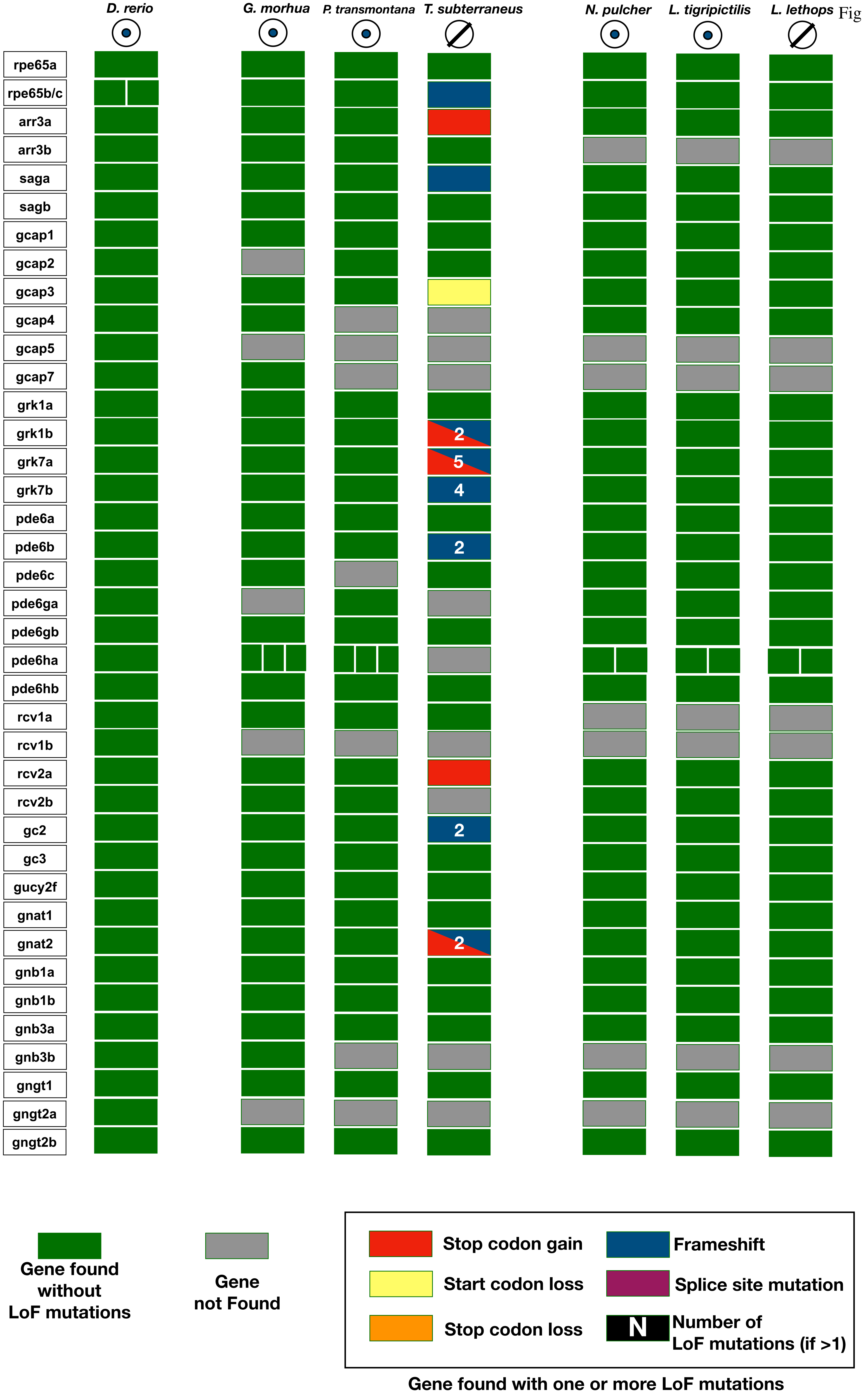
D. rerio
G. morhua
P. transmontana
T. subterraneus
N. pulcher
L. tigripictilis
L. lethops
rpe65a
rpe65b/c
arr3a
arr3b
saga
sagb
gcap1
gcap2
gcap3
gcap4
gcap5
gcap7
grk1a
grk1b
grk7a
grk7b
pde6a
pde6b
pde6c
pde6ga
pde6gb
pde6ha
pde6hb
rcv1a
rcv1b
rcv2a
rcv2b
gc2
gc3
gucy2f
gnat1
gnat2
gnb1a
gnb1b
gnb3a
gnb3b
gngt1
gngt2a
gngt2b
2
5
4
2
2
2
Gene found without LoF mutations
Gene not Found
Stop codon gain
Frameshift
Start codon loss
Splice site mutation
Stop codon loss
N
Number of LoF mutations (if >1)
Gene found with one or more LoF mutations

Fig. S2

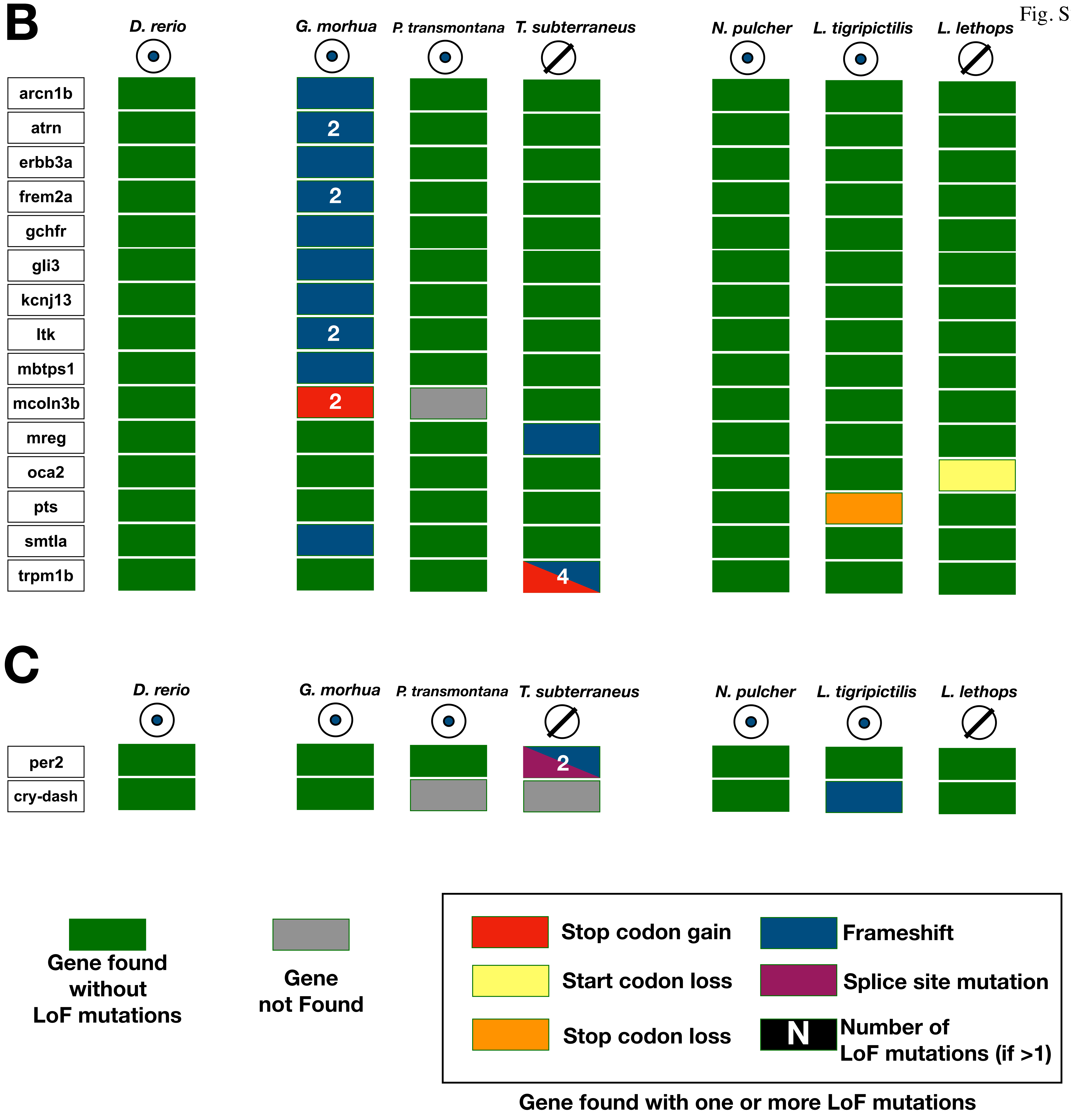
B
D. rerio
G. morhua
P. transmontana
T. subterraneus
N. pulcher
L. tigripictilis
L. lethops
arcn1b
atrn
erbb3a
frem2a
gchfr
gli3
kcnj13
ltk
mbtps1
mcoln3b
mreg
oca2
pts
smtla
trpm1b
2
2
2
2
4
C
D. rerio
G. morhua
P. transmontana
T. subterraneus
N. pulcher
L. tigripictilis
L. lethops
per2
cry-dash
2
Gene found without LoF mutations
Gene not Found
Stop codon gain
Frameshift
Start codon loss
Splice site mutation
Stop codon loss
N
Number of LoF mutations (if >1)
Gene found with one or more LoF mutations

Fig. S3

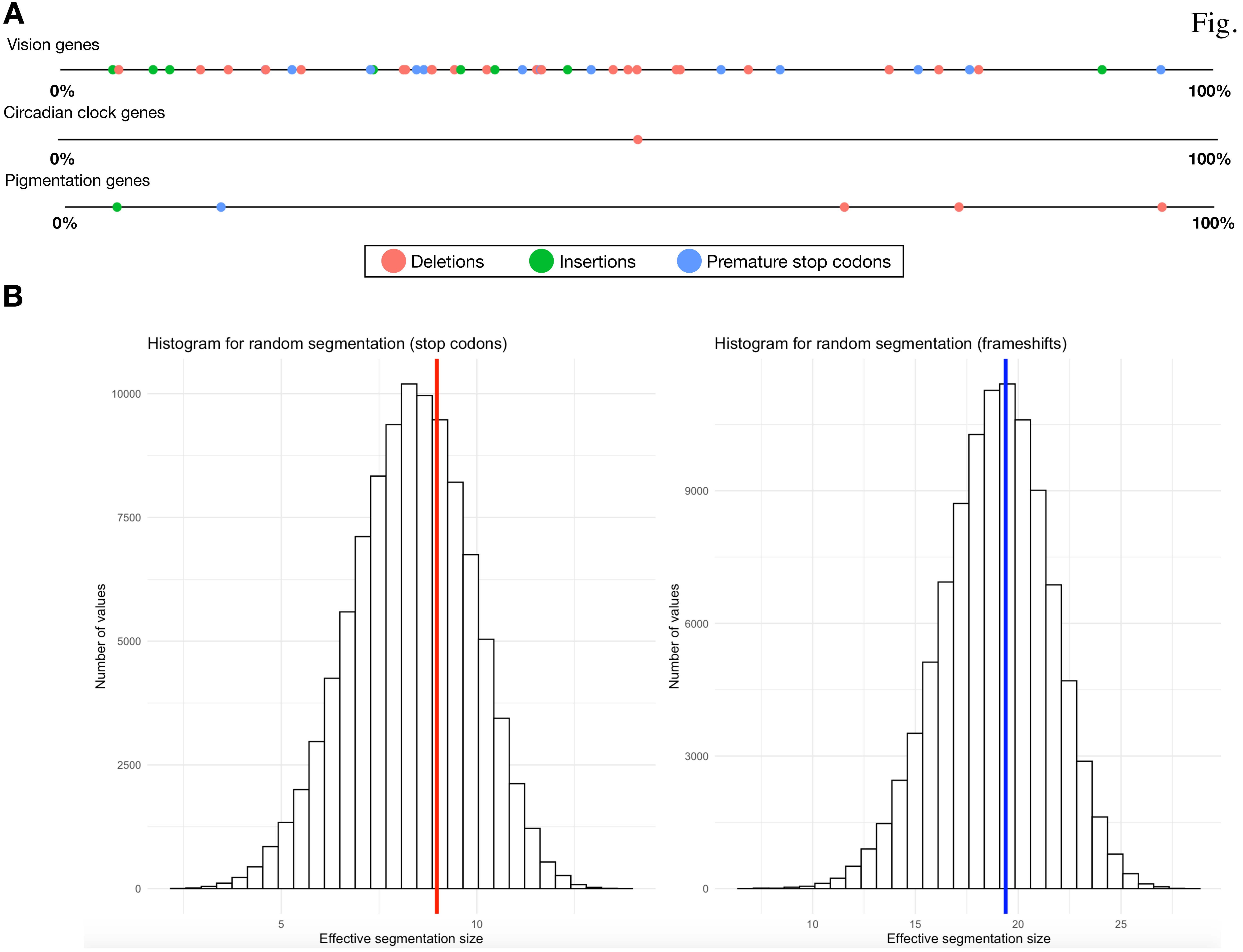
A
Vision genes
0%
100%
Circadian clock genes
0%
100%
Pigmentation genes
0%
100%
Deletions
Insertions
Premature stop codons
B
Histogram for random segmentation (stop codons)
Number of values
10000
7500
5000
2500
0
5
10
Effective segmentation size
Histogram for random segmentation (frameshifts)
Number of values
9000
6000
3000
0
10
15
20
25
Effective segmentation size

**A**

Fig. S4

### No gene is present on the region on which a LoF mutation is reported on Aaderma et al.

**Supplementary table S3 from Aaderma et al. 2020**

| Gene Symbol | Premature Stop Codon | Loss of Start Codon | Frameshift Mutation | Exon Splice Site Variant | Relevant GO Terms | Corresponding ZFIN ID | Implication in species with similar phenotypes | Specific Variant | Total Depth of Coverage ( number of reads with variant [percent of reads with variant], number of reads in forward direction, number of reads in reverse direction) | *Neolamprologous birchardi* Scaffold | *Neolamprologous birchardi* Scaffold Position of Variant | Exon effected by inactivating Mutation |
|---|---|---|---|---|---|---|---|---|---|---|---|---|
| pmch | X | | | | melanin-concentrating hormone activity | ZDB-GENE-041210-150 | None identified | C -> T | 79 (79 [100%], 43+, 36-) | NW_006272002.1 | 13048305 | 13048300-13048392 |
| oca2 | | X | | | melanin biosynthetic process, melanocyte differentiation, pigmentation | ZDB-GENE-070718-4 | Am: deletion (Protas et al. 2006, 2007; Klaassen et al. 2018) | T -> C | 54 (54 [100%], 27+, 27-) | NW_006272018.1 | 5745343 | 5745336-5745571 |
| opn4xa | X | | | | cellular response to light stimulus, phototransduction, visual perception | ZDB-GENE-110622-1 | None identified | C -> T | 76 (76 [100%], 26+, 50-) | NW_006272015.1 | 5259668 | 5259527-5259675 |
| rd3 | X | | | | retina development in camera-type eye* | ZDB-GENE-040724-103 | None (found to be expressed at similar levels in Am and Sa) | C->A | 52 (52 [100%], 25+, 27-) | NW_006272003.1 | 2415158 | 2414995-2415302 |
| rp1 | X | | | | photoreceptor cell development, retina development in camera-type eye | ZDB-GENE-120711-1 | Nonfunctional in subterranean mammals (Emerling and Springer 2014) | C -> T | 54 (54 [100%], 31+, 23-) | NW_006272040.1 | 3903778 | 3903601-3906062 |
| cry2 | X | | X | | response to light stimulus | ZDB-GENE-010426-6 | None identified | C->T (stop gained), Deletion; T (framshift) | 49 (49 [100%], 25+, 24-); 45 (45 ([100%], 23+, 22-) | NW_006272016.1 | 4605214 (stop gained), 4607152 (framshift) | 4604899-4605466, 4607131-4607356 |
| bfsp1 | X | | X | | lens fiber cell development† | ZDB-GENE-090608-2 | Upregulated in Sa (Meng et al. 2013) | C-> T (stop gained), Insertion; AGCAAGAAGT (framshift) | 57 (57 [100%], 33+, 24-) | NW_006272006.1 | 12555228 (stop gained), 12564282 (framshift) | 12554903-12555376, 12563304-12564856 |
| crybgx | | | X | | lens development in camera-type eye, visual perception | ZDB-GENE-070410-2 | Downregulated in Am (Hinaux et al. 2013) | Insertion; G | 50 (50 [100%], 23+, 27-) | NW_006272218.1 | 642367 | 642235-642461 |
| crybb3 | | | X | | lens development in camera-type eye, visual perception | ZDB-GENE-060130-2 | Premature stop codon in naked mole rat (Kim et al. 2011) | Deletion; TTGGGATTTGGGGTTGGGAGTAGGGTTAGGA | 12 (12 [100%], 5+, 7-) | NW_006272007.1 | 9006884 | 9006687-9007161 |
| crygm2f | | | X | | lens development in camera-type eye, visual perception | ZDB-GENE-080220-17 | Sa: upregulation (Meng et al. 2013) | Deletion; G | 54 (53 [98%], 31+, 23-) | NW_006272180.1 | 605286 | 605133-605375 |
| opn3 | | | X | | cellular response to light stimulus, phototransduction, response to stimulus, visual perception | ZDB-GENE-080227-16 | None identified | Deletion; C | 47 (46 [98%], 31+, 16-) | NW_006272062.1 | 3187434 | 3187203-3187499 |
| tmtops2a | | | X | | phototransduction, response to stimulus, visual perception | ZDB-GENE-130129-3 | Pa: premature stop codon (Cavallari et al. 2011) | Deletion; C | 49 (45 [92%], 33+, 16-) | NW_006272087.1 | 1747364 | 1747152-1747477 |
| impg2b | | | | X (segregating) | visual perception | ZDB-GENE-081031-56 | Am: Downregulated (Stahl and Gross 2017); Nonfunctional in subterranean mammals (Emerling and Springer 2014) | C -> A | 64 (30 [47%], 18+, 12-) | NW_006272014.1 | 2013719 | 2013689-2013721 |
| spx | X | | | | negative regulation of appetite | ZDB-GENE-041210-158 | None identified | C -> A | 62 (61 [98%], 28+, 33-) | NW_006272004.1 | 14758201 | 14758178-14758236 |
| ddb2 | | | X | | cellular response to DNA damage stimulus, DNA repair, response to UV | ZDB-GENE-050419-169 | Am: Constitutively expressed (Foulkes et al. 2016) | Deletion; G | 44 (44 [100%], 25+, 19-) | NW_006272016.1 | 1059370 | 1059282-1059437 |

**NCI genome viewer**

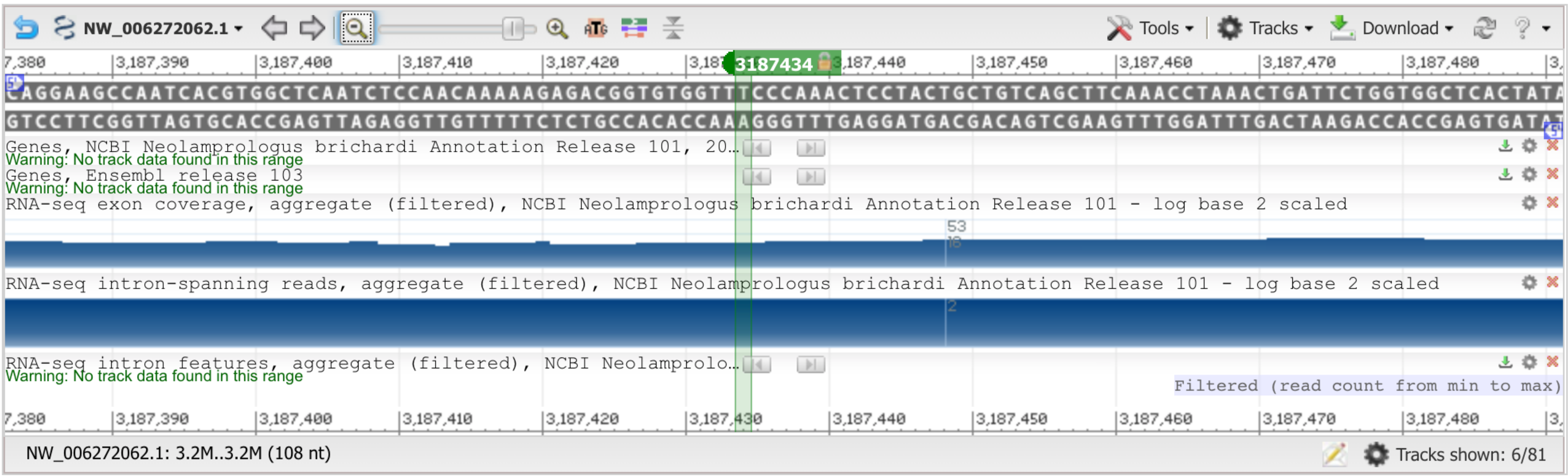

**B**

### Here is the exonerate result of opn3 mapped to the *N. brichardi* genome

Fig. S4

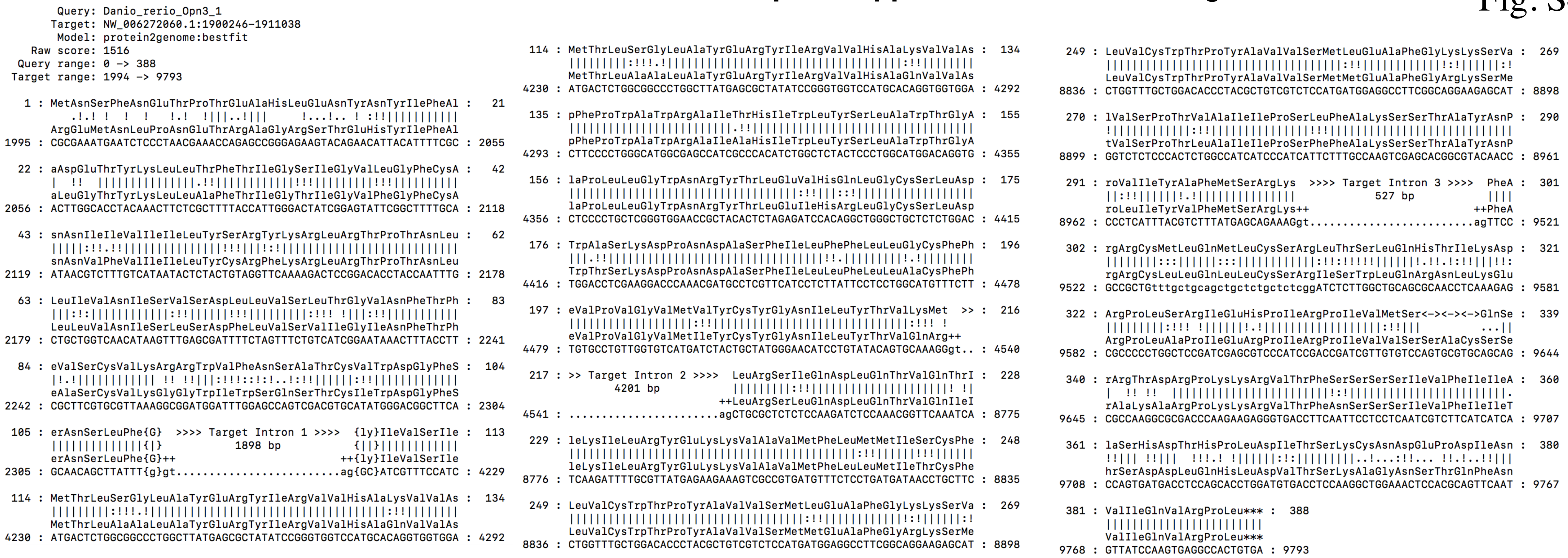


#### Here is the same region on the NCBI genome viewer :

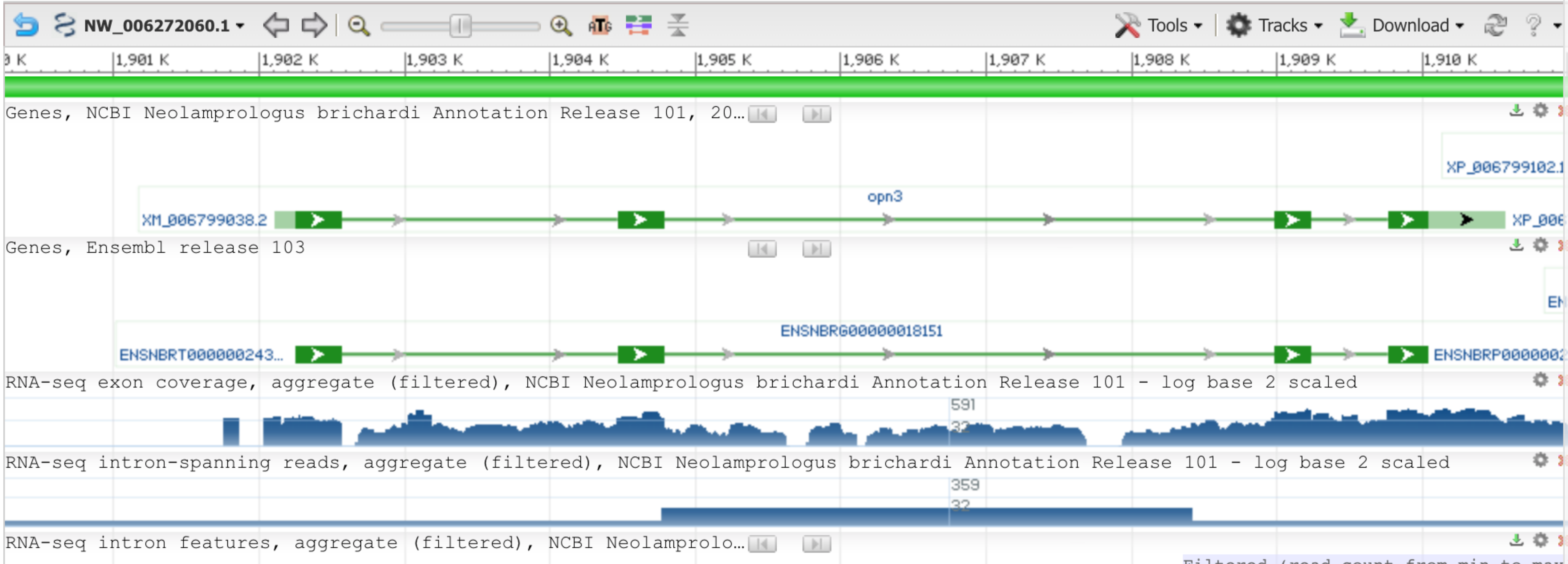

C

Fig. S4

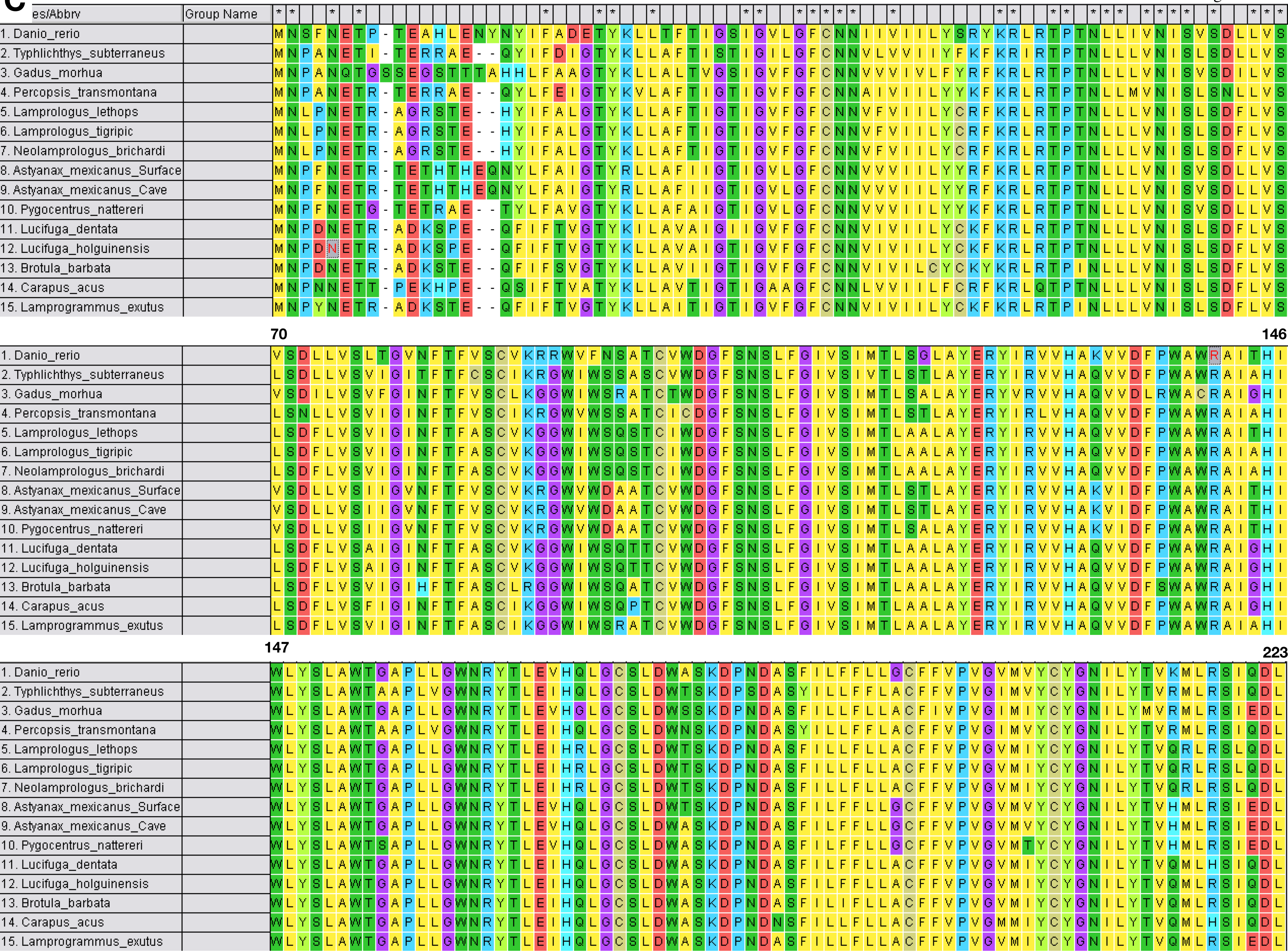

1
69
es/Abbrv
Group Name
1. Danio_rerio
2. Typhlichthys_subterraneus
3. Gadus_morhua
4. Percopsis_transmontana
5. Lamprologus_lethops
6. Lamprologus_tigripic
7. Neolamprologus_brichardi
8. Astyanax_mexicanus_Surface
9. Astyanax_mexicanus_Cave
10. Pygocentrus_nattereri
11. Lucifuga_dentata
12. Lucifuga_holguinensis
13. Brotula_barbata
14. Carapus_acus
15. Lamprogrammus_exutus
70
146
147
223

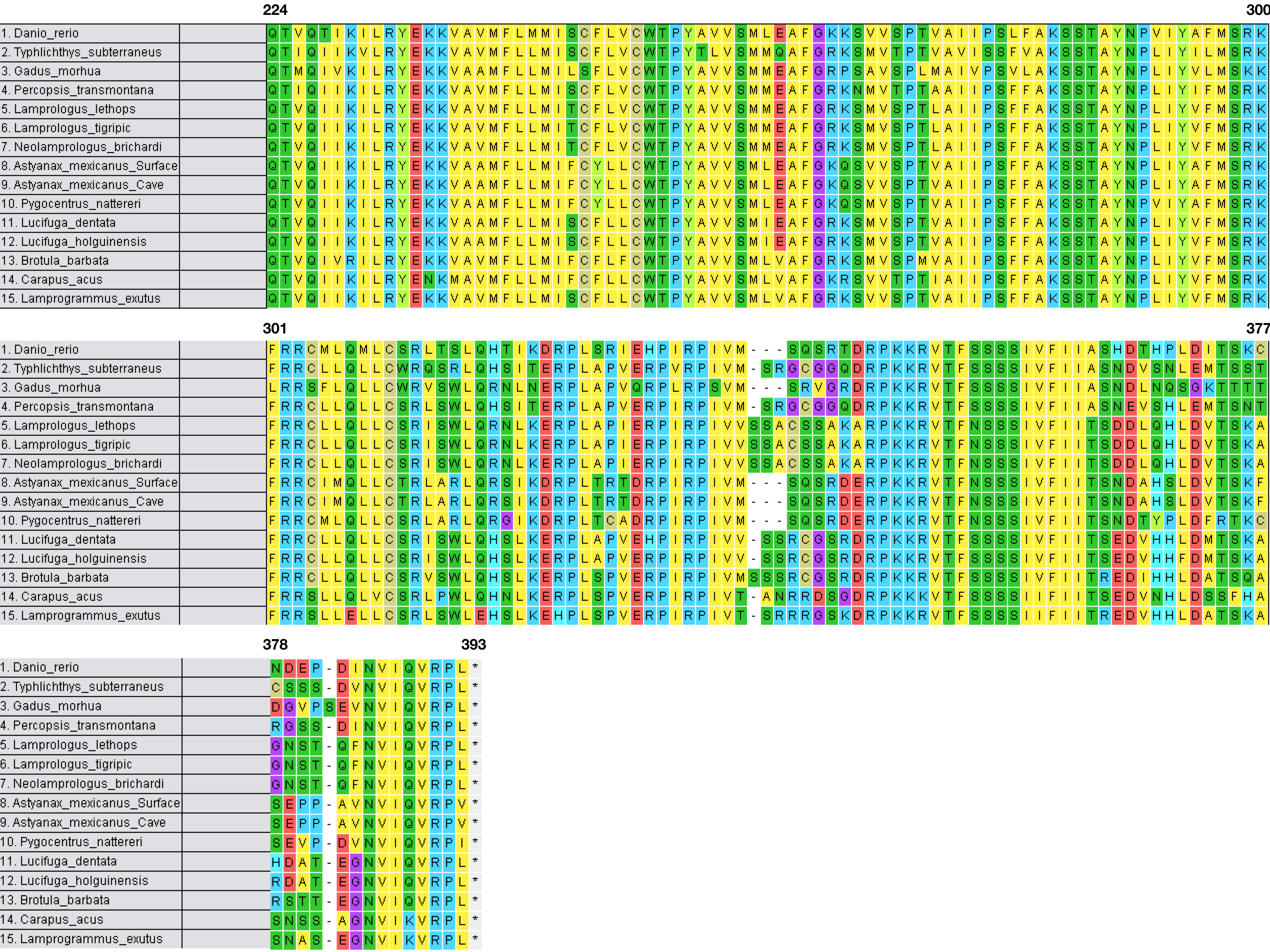
224
300
1. Danio_rerio QTVQTIKILRYEKKVAVMFLMMISCFLVCWTPYAVVSMLEAFGKKSVVSPTVAIIPSLFAKSSTAYNPVIYAFMSRK
2. Typhlichthys_subterraneus QTIQIIKVLRYEKKVAVMFLLMISCFLVCWTPYTLVSMMQAFGRKSMVTPTVAVISSFVAKSSTAYNPLIYILMSRK
3. Gadus_morhua QTMQIVKILRYEKKVAAMFLLMILSFLVCWTPYAVVSMMEAFGRPSAVSPLMAIVPSVLAKSSTAYNPLIYVLMSKK
4. Percopsis_transmontana QTIQIIKILRYEKKVAVMFLLMISCFLVCWTPYAVVSMMEAFGRKNMVTPTAAIIPSFFAKSSTAYNPLIYIFMSRK
5. Lamprologus_lethops QTVQIIKILRYEKKVAVMFLLMITCFLVCWTPYAVVSMMEAFGRKSMVSPTLAIIPSFFAKSSTAYNPLIYVFMSRK
6. Lamprologus_tigripic QTVQIIKILRYEKKVAVMFLLMITCFLVCWTPYAVVSMMEAFGRKSMVSPTLAIIPSFFAKSSTAYNPLIYVFMSRK
7. Neolamprologus_brichardi QTVQIIKILRYEKKVAVMFLLMITCFLVCWTPYAVVSMMEAFGRKSMVSPTLAIIPSFFAKSSTAYNPLIYVFMSRK
8. Astyanax_mexicanus_Surface QTVQIIKILRYEKKVAAMFLLMIFCYLLCWTPYAVVSMLEAFGKQSVVSPTVAIIPSFFAKSSTAYNPLIYAFMSRK
9. Astyanax_mexicanus_Cave QTVQIIKILRYEKKVAAMFLLMIFCYLLCWTPYAVVSMLEAFGKQSVVSPTVAIIPSFFAKSSTAYNPLIYAFMSRK
10. Pygocentrus_nattereri QTVQIIKILRYEKKVAAMFLLMIFCYLLCWTPYAVVSMLEAFGKQSMVSPTVAIIPSFFAKSSTAYNPVIYAFMSRK
11. Lucifuga_dentata QTVQIIKILRYEKKVAAMFLLMISCFLLCWTPYAVVSMIEAFGRKSMVSPTVAIIPSFFAKSSTAYNPLIYVFMSRK
12. Lucifuga_holguinensis QTVQIIKILRYEKKVAAMFLLMISCFLLCWTPYAVVSMIEAFGRKSMVSPTVAIIPSFFAKSSTAYNPLIYVFMSRK
13. Brotula_barbata QTVQIVRILRYEKKVAVMFLLMIFCFLFCWTPYAVVSMLVAFGRKSMVSPMVAIIPSFFAKSSTAYNPLIYVFMSRK
14. Carapus_acus QTVQIIKILRYENKMAVMFLLMIFCFLLCWTPYAVVSMLVAFGKRSVVTPTIAIIPSFFAKSSTAYNPLIYVFMSRK
15. Lamprogrammus_exutus QTVQIIKILRYEKKVAVMFLLMISCFLLCWTPYAVVSMLVAFGRKSVVSPTVAIIPSFFAKSSTAYNPLIYVFMSRK
301
377
1. Danio_rerio FRRCMLQMLCSRLTSLQHTIKDRPLSRIEHPIRPIVM---SQSRTDRPKKRVTFSSSSIVFIIASHDTHPLDITSKC
2. Typhlichthys_subterraneus FRRCLLQLLCWRQSRLQHSITERPLAPVERPVRPIVM-SRGCGGQDRPKKRVTFSSSSIVFIIASNDVSNLEMTSST
3. Gadus_morhua LRRSFLQLLCWRVSWLQRNLNERPLAPVQRPLRPSVM---SRVGRDRPKKRVTFSSSSIVFIIASNDLNQSGKTTTT
4. Percopsis_transmontana FRRCLLQLLCSRLSWLQHSITERPLAPVERPIRPIVM-SRGCGGQDRPKKRVTFSSSSIVFIIASNEVSHLEMTSNT
5. Lamprologus_lethops FRRCLLQLLCSRISWLQRNLKERPLAPIERPIRPIVVSSACSSAKARPKKRVTFNSSSIVFIITSDDLQHLDVTSKA
6. Lamprologus_tigripic FRRCLLQLLCSRISWLQRNLKERPLAPIERPIRPIVVSSACSSAKARPKKRVTFNSSSIVFIITSDDLQHLDVTSKA
7. Neolamprologus_brichardi FRRCLLQLLCSRISWLQRNLKERPLAPIERPIRPIVVSSACSSAKARPKKRVTFNSSSIVFIITSDDLQHLDVTSKA
8. Astyanax_mexicanus_Surface FRRCIMQLLCTRLARLQRSIKDRPLTRTDRPIRPIVM---SQSRDERPKKRVTFNSSSIVFIITSNDAHSLDVTSKF
9. Astyanax_mexicanus_Cave FRRCIMQLLCTRLARLQRSIKDRPLTRTDRPIRPIVM---SQSRDERPKKRVTFNSSSIVFIITSNDAHSLDVTSKF
10. Pygocentrus_nattereri FRRCMLQLLCSRLARLQRGIKDRPLTCADRPIRPIVM---SQSRDERPKKRVTFNSSSIVFIITSNDTYPLDFRTKC
11. Lucifuga_dentata FRRCLLQLLCSRISWLQHSLKERPLAPVEHPIRPIVV-SSRCGSRDRPKKRVTFSSSSIVFIITSEDVHHLDMTSKA
12. Lucifuga_holguinensis FRRCLLQLLCSRISWLQHSLKERPLAPVERPIRPIVV-SSRCGSRDRPKKRVTFSSSSIVFIITSEDVHHFDMTSKA
13. Brotula_barbata FRRCLLQLLCSRVSWLQHSLKERPLSPVERPIRPIVMSSSRCGSRDRPKKRVTFSSSSIVFIITREDIHHLDATSQA
14. Carapus_acus FRRSLLQLVCSRLPWLQHNLKERPLSPVERPIRPIVT-ANRRDSGDRPKKKVTFSSSSIIFIITSEDVNHLDSSFHA
15. Lamprogrammus_exutus FRRSLLELLCSRLSWLEHSLKEHPLSPVERPIRPIVT-SRRRGSKDRPKKRVTFSSSSIIFIITREDVHHLDATSKA
378
393
1. Danio_rerio NDEP-DINVIQVRPL*
2. Typhlichthys_subterraneus CSSS-DVNVIQVRPL*
3. Gadus_morhua DGVPSEVNVIQVRPL*
4. Percopsis_transmontana RGSS-DINVIQVRPL*
5. Lamprologus_lethops GNST-QFNVIQVRPL*
6. Lamprologus_tigripic GNST-QFNVIQVRPL*
7. Neolamprologus_brichardi GNST-QFNVIQVRPL*
8. Astyanax_mexicanus_Surface SEPP-AVNVIQVRPV*
9. Astyanax_mexicanus_Cave SEPP-AVNVIQVRPV*
10. Pygocentrus_nattereri SEVP-DVNVIQVRPI*
11. Lucifuga_dentata HDAT-EGNVIQVRPL*
12. Lucifuga_holguinensis RDAT-EGNVIQVRPL*
13. Brotula_barbata RSTT-EGNVIQVRPL*
14. Carapus_acus SNSS-AGNVIKVRPL*
15. Lamprogrammus_exutus SNAS-EGNVIKVRPL*

Fig. S5

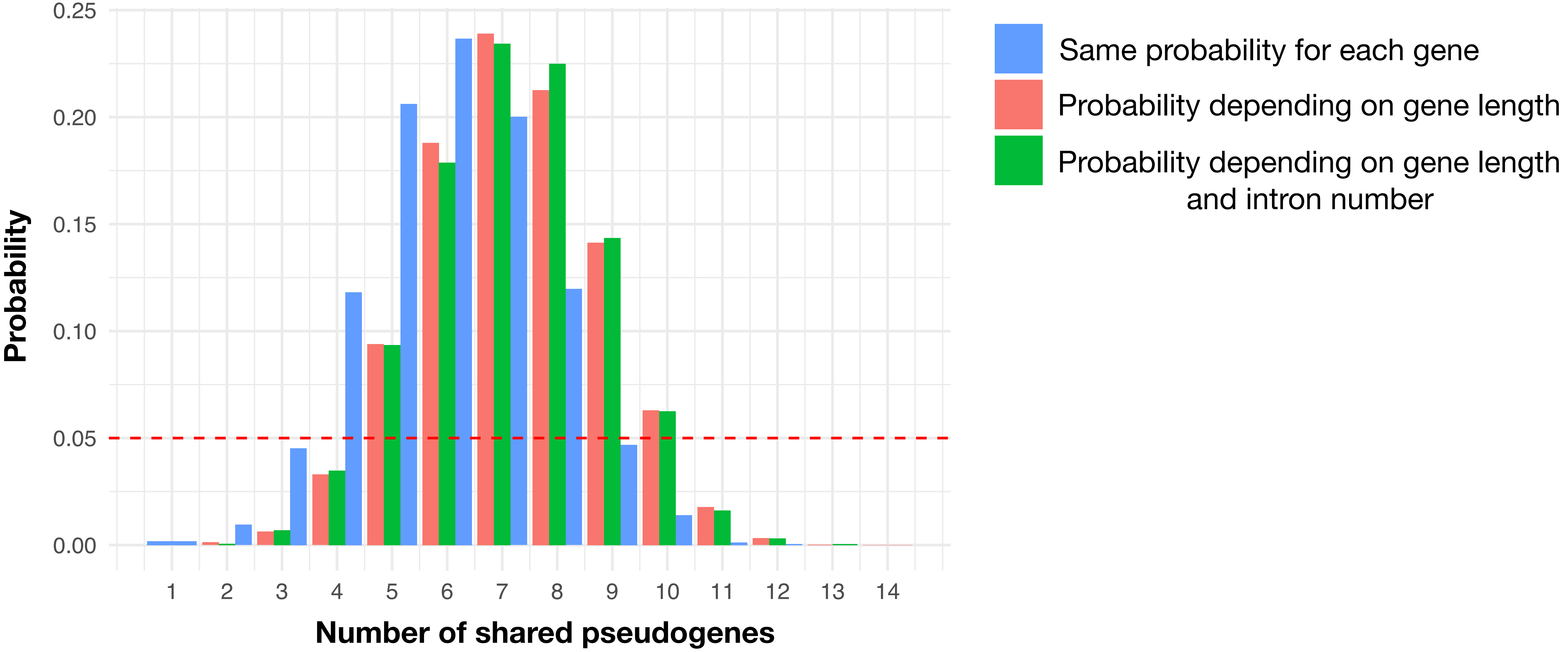

0.25
0.20
0.15
0.10
0.05
0.00
Probability
1
2
3
4
5
6
7
8
9
10
11
12
13
14
Number of shared pseudogenes
Same probability for each gene
Probability depending on gene length
Probability depending on gene length and intron number
